# Examination of zinc in the circadian system

**DOI:** 10.1101/790352

**Authors:** Mahtab Moshirpour, Amy S. Nakashima, Nicole Sehn, Victoria M. Smith, Richard H. Dyck, Michael C. Antle

**Affiliations:** Department of Psychology, University of Calgary, Calgary, Alberta, Canada; Hotchkiss Brain Institute, University of Calgary, Calgary, Alberta, Canada; Department of Cell Biology and Anatomy, University of Calgary, Calgary, Alberta, Canada; Department of Physiology & Pharmacology, University of Calgary, Calgary, Alberta, Canada

**Keywords:** Zn^2+^, rhythms, photic, c-fos, mper1, ipRGCs

## Abstract

Zinc is a trace element that is essential for a large number of biological and biochemical processes in the body. In the nervous system zinc is packaged into synaptic vesicles by the ZnT3 transporter, and synaptic release of zinc can influence the activity of postsynaptic cells, either directly though its own cognate receptors, or indirectly by modulating activation of receptors for other neurotransmitters. Here, we explore the anatomical and functional aspects of zinc in the circadian system. Melanopsin-containing retinal ganglion cells in the mouse retina were found to colocalize ZnT3, indicating that they can release zinc at their synaptic targets. While the master circadian clock in the hamster suprachiasmatic nucleus (SCN) was found to contain, at best, sparse zincergic input, the intergeniculate leaflet (IGL) was found to have prominent zincergic input. Levels of zinc in these areas were not affected by time of day. Additionally, IGL zinc staining persisted following enucleation, indicating other prominent sources of zinc instead of, or in addition to, the retina. Neither enhancement nor chelation of free zinc at either the SCN or IGL altered circadian responses to phase-shifting light in hamsters. Finally, entrainment, free-running, and circadian responses to light were explored in mice lacking the ZnT3 gene. In every aspect explored, the ZnT3-KO mice were not significantly different from their wildtype counterparts. These findings highlight the presence of zinc in areas critical for circadian functioning but have yet to identify a role for zinc in these areas.

**Highlights:**

- The synaptic zinc transporter ZnT3 is found in melanopsin-containing retinal ganglion cells.
- While zinc input to the hamster SCN was found to be sparse at best, prominent zincergic staining was found throughout the IGL.
- Zinc levels in the SCN and IGL did not change between the night and day.
- Neither increasing nor decreasing zinc levels in either the SCN or IGL had an influence on circadian responses to light.
- Mice lacking the ZnT3 transporter did not differ from wildtype mice on a wide variety of circadian measures.

## INTRODUCTION

Daily rhythms in physiology and behavior are regulated in mammals by the master circadian clock located in the suprachiasmatic nucleus (SCN;Antle and Silver, 2005). This clock receives input from the retina so that the rhythms it generates are coordinated with the light/dark cycle of the environment (Rusak and Zucker, 1979). Specialized intrinsically photosensitive retinal ganglion cells (ipRGCs) that use the photopigment melanopsin provide information about environmental light to the circadian system (Berson et al., 2002;Hattar et al., 2002;Moore et al., 1995;Provencio et al., 2000). Axons from these ipRGCs give rise to the retinohypothalamic tract that innervates the ventrolateral region of the SCN, and uses glutamate as its primary neurotransmitter (Ebling, 1996). The circadian clock also receives input from other brain regions, and these inputs can modulate how the circadian clock responds to light and glutamate. Serotonin from the raphe nuclei attenuates responses to light and glutamate (Antle et al., 2003;Mistlberger and Antle, 1998;Rea et al., 1994). The retinorecipient intergeniculate leaflet (IGL) provides NPY-ergic innervation to the SCN that also opposes circadian responses to light (Biello et al., 1997). Identifying other such modulatory substances could help in treating sleep and circadian disorders. One such candidate is the divalent cation zinc.

The signaling pool of zinc is found in the brain, stored in vesicles of glutamatergic neurons in the cortex, amygdala and hippocampus (Brown and Dyck, 2004) from which it is released and can act as a co-transmitter and neuromodulator at post-synaptic terminals. Zinc is packaged into vesicles exclusively through the ZnT3 transporter (slc30a3;Cole et al., 1999;McAllister and Dyck, 2017;Nakashima and Dyck, 2009;Palmiter et al., 1996). Vesicular zinc inhibits activation of NMDA receptors that contain the GluN2A subunit (Paoletti et al., 1997;Paoletti et al., 2009), while it facilitates activation of non-NMDA receptors (reviewed in McAllister and Dyck, 2017;Ugarte and Osborne, 2001). GABA_A_ receptors are also modulated by zinc (Huang et al., 1993;Kawahara et al., 1993;Strecker et al., 1999). Zinc has also been shown to be the primary endogenous ligand for GPR39, a metabotropic receptor expressed in the brain that signals through the Gq-PLC-IP3 pathway (Besser et al., 2009).

The possibility that zinc can modulate circadian rhythms is suggested by its presence in the ganglion cell layer of the retina (Redenti and Chappell, 2004). Zinc has been histochemically located in the ventrolateral portion of the rat SCN (Huang, et al., 1993). In dissociated SCN neurons, zinc enhances the potassium I_A_ current (Huang, et al., 1993), inhibits GABA_A_ currents (Kawahara, et al., 1993;Strecker, et al., 1999), and inhibits NMDA GluNR2A currents (Clark and Kofuji, 2010). These in vitro findings suggest that zinc could modulate circadian responses. To explore this possibility more fully, we devised a series of anatomical and behavioral studies to determine if vesicular zinc might modulate circadian rhythms. Using hamsters and mice, we explore the localization of vesicular zinc and its transporter ZnT3 in the retina and circadian network. We explore the effects that chelating or enhancing zinc levels in the SCN and IGL has on circadian responses to light. Finally, we explore the circadian properties of the ZnT3-knockout (KO) mouse.

## EXPERIMENTAL PROCEDURES

### Animals and housing

Male C57BL/6J wildtype adult mice (n=4) from an in-house colony were used for retinal ZnT3 and melanopsin immunohistochemistry. Syrian hamsters (n=40) obtained from Charles River Laboratories (Kingston, NY, USA) were used for sodium selenite injections and the vesicular zinc staining, enucleation, and photic entrainment experiments (detailed below). ZnT3-KO mice (n=9; B6;129-*Slc30a3*^*tm1Rpa*^) and wildtype littermate controls (n=9) were bred from heterozygous animals from an in-house colony founded by animals received from Dr. Richard Palmiter, University of Washington. ZnT3-KO mice were genotyped using the protocol for this strain provided by The Jackson Laboratory (https://www.jax.org/strain/005064) as previously described (Cole et al., 1999;Nakashima et al., 2011).

Animals were initially housed in pairs in a temperature and humidity-controlled room with either a 14:10 (hamsters) or 12:12 (mice) light-dark cycle (LD; approximately 1500 lux during the light phase) and provided with food and water *ad libitum*. Animals used in behavioural experiments were transferred to individual polycarbonate cages equipped with a running wheel. Periodic cage changes took place seven days prior and following manipulation days. The wheel running activity of the animals was recorded by means of a magnetic switch fastened to each wheel and monitored using Clocklab (Actimetrics, Wilmette, IL). All experiments adhered to the guidelines of the Canadian Council on Animal Care and were approved by the Life and Environmental Sciences Animal Care Committee at the University of Calgary.

### Surgery

Some hamsters were surgically implanted with cannulas aimed at the SCN (unilateral, n=12) or the IGL (bilateral, n=14). Each animal received a subcutaneous injection of the analgesic butorphanol (2 mg/kg; Wyeth, Madison, NJ, USA) and an i.p. injection of the anesthetic sodium pentobarbital (90 mg/kg; CEVA). While under anesthesia, hamsters were stereotaxically implanted with either one (SCN) or two (IGL) 9 mm 22-gauge stainless steel cannulae (Plastics One Inc., Roanoke, VA, USA). The cannulae were cemented to the skull using dental acrylic and jeweler’s screws. Coordinates for the SCN were 0.0 mm anterior to bregma, 0.3 mm lateral to the midline, 7.0 mm ventral to the skull surface, and coordinates for the IGL were 1.9 mm posterior to bregma and 3.3 mm lateral to the midline. To determine depth, two targets were calculated at 3.3 mm and 4.8 mm below dura and the skull, respectively. The average of the two target values was taken as the dorsoventral coordinate. The incisor bar was set to 2 mm below the interaural level for all surgeries. A dummy cannula was inserted to maintain patency.

Brains were collected for histology at the end of the experiment using the perfusion method described above. Brain sections were stained using cresyl violet and coverslipped in Permount. Animals were excluded if the tip of the cannula was more than 600μm away from the margin of the SCN, or if the tip of either one of the two cannulae was more than 600 μm away from the margin of either IGL.

Some hamsters received bilateral enucleations. Hamsters were anesthetized with sodium pentobarbital (100-120 mg/kg) and provided with butorphanol as a pre-operative analgesic and given a subcutaneous injection of 0.5% bupivacaine into the area immediately around eye. When unconscious, the eye was gently dissected, and the optic nerve severed. Post-operative analgesic (butorphanol) was administered as needed.

### Histology

#### Autometallographic Zinc Staining

The autometallographic zinc staining procedure was adapted from the technique previously employed by (Danscher, 1982). Hamsters received an i.p. injection of 15mg/kg of sodium selenite dissolved in molecular grade water in DD conditions with the aid of night vision goggles (BG15Alista, Richmond Hill, Ontario, Canada) at either ZT6 or ZT18. One hour later, animals were euthanized with an overdose of sodium pentobarbital (250mg/kg Euthanyl). For experiments where the tissue was only being processed for autometallographic zinc staining, brains were rapidly extracted and immediately frozen on dry ice. If alternate sections were being processed using immunohistochemistry, animals were perfused transcardially with ∼50ml of cold phosphate buffered saline (PBS) followed by 50ml of cold 4% paraformaldehyde (PFA) in PBS. Frozen brain sections (35 µm) were collected through the SCN and IGL with a cryostat and were mounted directly onto gelatin-coated microscope slides.

The tissue was allowed to thaw at room temperature and was then rehydrated in an alcohol series (95% EtOH for 15 min, 70% for 2 min, 50% for 2 min, 3×2 min in dH2O). Slides were then dipped in gelatin and transferred to a silver lactate solution for physical development. The developer solution was prepared by mixing the following solutions in order: 50% gum arabic from African Acacia trees (100 g in 200 ml dH2O), citrate buffer (5.1g citric acid, 4.7 g sodium citrate, 20 ml dH2O), silver lactate (0.22g in 30 ml dH2O), and hydroquinone (1.7g in 20 ml dH2O). Slides were developed for 120-150 min in the dark and were checked every 10-15 min, starting at 100 min. Slides were removed from the developer solution after the tissue had obtained a dark brown uniform stain. The slides were then gently washed under slowly running tap water at 37°C for 10 min and then in dH2O (2×3 min). Slides were then exposed to 5% sodium thiosulfate in dH2O for 12 min and washed in dH2O (2×2 min). Sections underwent a final alcohol dehydration series, were cleared with xylene and coverslipped with Permount.

#### Immunohistochemistry

Immunohistochemistry protocols were adapted from those previously used in our lab (Smith et al., 2010). Animals received an overdose of sodium pentobarbital (Euthanyl, 250mg/kg) and were then transcardially perfused with cold PBS followed by cold 4% PFA in PBS. Depending on the experiment, brains or eyes were extracted and were post-fixed overnight at 4°C in 4% PFA followed by 24h in 20% sucrose in PBS. Frozen sagittal sections (20µm for eyes, 35µm for brains) were collected with a cryostat and were mounted on gelatin-coated slides (eyes) or were collected into PBS baths (brains). The general immunohistochemistry procedure involved three 10 min washes in 0.3% PBS with Triton-X-100 (PBSx), followed by a 60-min incubation in blocking buffer (10% normal donkey serum for immunofluorescence, 10% normal goat serum for diaminobenzidine (DAB) staining). Following this, the tissue was incubated in the primary antibodies for 48h at 4°C. Tissue was then rinsed again (six 10-min washes) before being incubated in the secondary antibodies. For tissue to be stained with DAB, the tissue was then incubated in an avidin-biotin complex bath (Vectastain elite, Vector Laboratories, Burlingame, CA). The tissue was then developed for approximately 5 min with 0.05 % DAB and 0.02 % NiCl in 0.1 M Tris buffer activated with 80 μL of 30 % H_2_O_2_. Brain slices were dehydrated through an alcohol series, cleared with xylenes and coverslipped with Permount (Fisher Scientific, Ottawa, ON, USA) for DAB tissue, or with Krystalon for immunofluorescent tissue.

Retinal immunohistochemistry included a 20 min wash in 4% PFA prior to initiation of the protocol detailed above. Primary antibodies used were for melanopsin (anti-rabbit 1:5000, Advanced Targeting Systems, San Diego, CA, USA) and ZnT3 (anti-mouse 1:800, Synaptic Systems, Goettingren, Germany). Secondary antibodies were CY-3 donkey anti-rabbit and CY-2 donkey anti-mouse (1:200, Jackson ImmunoResearch Laboratories Inc. West Grove, PA, USA). Neuropeptide Y immunohistochemistry included a 30 min incubation in 0.5% H_2_O_2_ prior to initiation of the protocol detailed above. The primary antibody was rabbit anti-NPY (1:10,000; ImmunoStar, Hudson, WI, USA), and a goat-anti rabbit biotinylated secondary was employed (1:200; Vector Laboratories, Burlingame, CA, USA).

Mouse SCN immunohistochemistry used the follow primary antibodies: Fos (goat anti-Fos; 1:20,000; Santa Cruz Biotechnology, Santa Cruz, CA, USA), vasopressin (VP; guinea-pig anti-VP; 1:5000; Peninsula Laboratories, San Carlos, CA, USA), and bombesin/gastrin-releasing peptide (GRP; rabbit anti-GRP; 1:5000; ImmunoStar Inc., Hudson, WI, USA). The secondary antibodies used were CY-2 donkey anti-rabbit, CY-3 donkey anti-goat, and CY-5 donkey anti-guinea-pig (all 1:200; Jackson ImmunoResearch Laboratories Inc., West Grove, PA, USA).

#### In situ hybridization

In situ hybridization followed our previously described protocol (Smith et al., 2008). Briefly, tissue was first treated with proteinase K (1mg/ml, 0.1 M Tris buffer pH 8.0, 50 mM EDTA; 10 min) at 37°C. This reaction was terminated with the addition of 4% PFA. Tissue was rinsed in saline sodium citrate (300 mM NaCl, 30 mM sodium citrate), and then treated with 0.25% acetic anhydride in 0.1M triethanolamine for 10 min. The tissue was then incubated in 1.5 ml hybridization buffer (50% formamide, 60 mM sodium citrate, 600 mM NaCl, 10% dextran sulphate, 1% N-laurylsarcosine, 25 mg/ml tRNA, 1x Denhardt’s, 0.25 mg/ml salmon sperm DNA) which also contained the DIG-labelled sense and antisense *mPer1* (mPer1 plasmid generously provided by Dr. L. Yan, Michigan State University, East Lansing, MI, USA) probes for 16 h at 60°C. Following high-stringency post-hybridization washes (50% formamide/saline sodium citrate at 60°C, 2×30 min), sections were treated with RNaseA, and then underwent more high-stringency washes. Tissue was then processed for immunodetection using a nucleic acid detection kit (Roche, Indianapolis, IN, USA). After incubation for 1 h at room temperature in 1.0% of blocking reagent in buffer one (100mM Tris-HCL buffer, 150 mM NaCl, pH 7.5), sections were incubated in alkaline phosphatase-conjugated DIG antibodies diluted in buffer 1(1:5000) for 3 days at 4°C. Sections were then washed in buffer 1 (2×15 min) and incubated in buffer 3 (100 mM Tris-HCL buffer, pH 9.5, containing 100 mM NaCl and 50 mM MgCl_2_) for 5 min and then incubated in a solution containing nitroblue tetrazolium salt (0.34 mg/ml) and 5-bromo-4-chloro-3-indolyl phosphate toluidine salt (0.18 mg/ml) for 16 h. The staining reaction was stopped by placing the sections in buffer 4 (10 mM Tris-HCL containing 1 mM EDTA, pH 8.0). Tissue was then mounted on gelatin coated slides, air dried, dehydrated in an ascending alcohol series, cleared in xylene, air dried, and coverslipped with Permount.

#### Image Analysis

All image analysis was performed on an Olympus BX51 microscope equipped for both brightfield and epifluorescence. Images were captured with a QI CAM Fast 1394 cooled CCD camera (QImaging, Burnaby, BC, Canada) connected to a computer running Image-Pro Plus software (Media Cybernetics, Inc. Rockville, MD).

##### Retinal histology

In total, 120 cells were identified. Colocalization was determined by initially identifying a melanopsin cell in the retinal ganglion cell layer (on the CY-3 filter channel) by its characteristic shape and extensive dendritic networks. The presence or absence of ZnT3 protein was then assessed. The signals were considered colocalized if the ZnT3 signal was of the same shape, size, and position as the initial melanopsin signal.

##### Zinc Histology

All images were captured with the same lighting conditions and exposure time. Densitometric analysis was completed on 16-bit images with the use of ImageJ software (ImageJ 1.42q; National Institutes of Health, Bethesda, MD). Zinc staining was quantified for the SCN by measuring the density of staining in each SCN nucleus and comparing the values to the density in the optic chiasm. The relative optical density (ROD) was calculated by taking the ratio of the SCN to optic chiasm density for the dorsolateral, dorsomedial and ventral regions. Density values were averaged across the SCN nuclei and ROD values were averaged across three consecutive SCN sections. ROD of IGL sections was determined by taking the ratio of staining in the IGL to staining in the dorsolateral geniculate nucleus (dLGN) for the left and right IGL. Representative rostral, mid and caudal IGL sections were averaged for the left and right IGL. Statistical significance was set at *p*<0.05 for all tests. Separate two-tailed independent samples t-tests were conducted to analyze the ROD between the two time points for the dorsomedial, dorsolateral and ventral SCN, and the left and right IGL.

##### SCN histology

Tissues triple labelled for cFos, AVP, and GRP were captured. cFos-immunoreactive (-IR) cells were counted bilaterally at the mid-caudal level of the SCN. Cell counts were obtained in the SCN shell (delineated by AVP-IR cells) and in the SCN core (delineated by GRP-IR cells). Counts were conducted by two individuals blind to the treatment conditions. Images of DIG-labelled cells were captured using a RGB colour filter using the microscope described above. Measurements of ROD of staining were obtained bilaterally at the mid-caudal level of the SCN using ImageJ (ImageJ 1.34s; National Institutes of Health, Bethesda, MD, USA). The core and shell were analyzed separately by two individuals blind to the treatment conditions.

### Experiment 1- Confirming ZnT3 localization in ipRGCs

We first explored if the ZnT3 zinc transporter was found in melanopsin-IR ganglion cells in the mouse retina. Mice (n=4) were euthanized, perfused and their eyes were collected for double-label fluorescent immunohistochemistry as described above. Mouse tissue was used to explore this question as these antibodies did not produce specific staining in hamster retinas in our hands. We identified 120 melanopsin-IR cells. These were determined to be ZnT3-IR if signal on the ZnT3 channel had the same size, shape and location as the soma on the melanopsin channel.

### Experiment 2- Histochemical zinc staining

Eight male Syrian hamsters were used for visualization of histochemical zinc in the SCN and IGL. Two weeks prior to the beginning of the experiment, the animals were transferred to individual cages and kept under the same light-dark cycle. Two days prior to receiving sodium selenite injections, all animals were moved to constant dark (DD). Zinc terminal staining was visualized at two specific time points: the middle of the former light period (defined as zeitgeber time, ZT6) and the middle of the former dark period (ZT18), where lights off in the former LD cycle is designated ZT12 by convention (n=4 per group).

### Experiment 3- Does zinc input to the circadian system come from the retina?

To test the hypothesis that the retina is the source of zinc input to the circadian system, six male hamsters (controls n=3, n=3 experimental animals) underwent a bilateral enucleation surgery as described above. As experiment #2 revealed prominent staining in the IGL but only sparse staining in the SCN, we focused on the IGL in this experiment. A week of recovery was allowed to provide time for the degeneration of remaining retinal projections. Hamsters then received an i.p. injection of sodium selenite (15mg/kg). An hour later, animals were euthanized with an overdose of sodium pentobarbital (250mg/kg) and were then perfused transcardially with ∼50ml of cold PBS followed by 50ml of cold 4% PFA in PBS. Brains were PFA fixed in this experiment to allow alternate sections to be stained for NPY to delineate the IGL. Sections were collected as described above, with one series being processed for zinc as described above. The alternate series was immunostained for NPY. Relative optical density was used to assess density of zincergic terminals in the IGL of both control and enucleated hamsters.

### Experiment 4- Effect of zinc level modulation on circadian responses to light

To test the effects of manipulating zinc levels on circadian responses to light, male hamsters underwent cannula implantation surgery (SCN, n=12; IGL, n=14) as described above. Zinc chloride (ZnCl_2_; Sigma) dissolved in 0.9% saline to a final concentration of 50μM was used to increase extracellular zinc (Takeda et al., 2003), while N,N,N′,N′-Tetrakis(2-pyridylmethyl)ethylenediamine (TPEN; Sigma) dissolved in 10% dimethylsulfoxide (DMSO) to a final concentration of 5mM was used as a zinc chelator (Cuajungco and Lees, 1996). Vehicle controls of 0.9% sterile saline and 10% DMSO were included for the zinc donor and chelator, respectively. Drugs were administered to animals based on a counterbalanced design throughout the course of the experiment. Due to deterioration of rhythmicity with repeated SCN injections, only one vehicle control manipulation was performed on each animal (either saline or 10%DMSO). Repeated IGL injections did not lead to the same deterioration of rhythmicity, so these animals received both vehicle treatments in addition to the ZnCl_2_ and TPEN treatments. Intracranial injections of 0.5μL of drugs were administered over a period of 30 seconds during each manipulation by means of a 26-gauge injection cannula attached to a 1μL Hamilton syringe with polyethylene 20 tubing. The injection cannula extended 1mm beyond the tip of the implanted guide cannula. All injections were made in DD conditions with the aid of night-vision goggles. Ten minutes following drug administration, animals were placed in light boxes and were exposed to a 15-min 40 lux light pulse in the late subjective night (circadian time (CT18, 6 h after activity onset, which is defined as CT12 by convention). Phase shifts were analyzed with Clocklab by fitting a regression line to activity onsets for the 10 days prior to the manipulation and another to onsets on days 3-10 following the manipulation. The horizontal difference between these lines on the day following the manipulation was used as the phase shift.

All comparisons were made using SigmaPlot (Systat Software, Inc.; San Jose, CA). Statistical significance was set at *p*<0.05 for all tests. A one-way repeated measures ANOVA was used to determine whether there was a statistically significant difference in the photic phase shifts between the conditions (zinc donor, zinc chelator, vehicle control). For the SCN injections, animals only received one of the two vehicles (either saline or DMSO). A two-tailed independent samples t-test was used to compare phase shifts in these vehicle conditions, and they were merged for the sake of the ANOVA analysis if there was no difference between the vehicle treatment. For the IGL treatments, animals received both vehicle treatments. A two-tailed paired t-test was used to compare these shifts, and as they did not significantly differ, the average of these was used for the ANOVA analysis. All means are reported as ± standard error of the mean (SEM) in the figures and as ±standard deviation in the text.

### Experiment 5- Circadian rhythms changes in the ZnT3 KO model

#### Circadian Behavior

To explore the consequences of global loss of vesicular zinc on circadian rhythms, we used mice lacking ZnT3. These mice exhibit a complete lack of vesicular zinc in the brain (Cole, et al., 1999;McAllister and Dyck, 2017). Mice were initially housed in an LD cycle. After two weeks in LD, the mice underwent a masking paradigm, whereby for 24 h, the LD cycle was 2:2. Mice were then returned to LD for two more weeks before being transferred to DD. After two weeks in DD, the mice were exposed to a dim light pulse (15 min, 40 lux) at either CT16 (early night) or CT22 (late night). Two weeks later, the mice were exposed to a light pulse again, however this time undergoing the pulse at the other time. Following this, the same paradigm of light pulses was repeated; however, in the second round, bright light (∼1200 lux) was used. Two weeks following this manipulation, animals were exposed to a bright light pulse (∼1200 lux) during the subjective day (CT6).

#### Circadian analysis

Basic features of the circadian rhythms of the ZnT3-KO mice were quantified using mice running wheel data collected in Clocklab as previously described (Smith et al., 2015). The LD ratio, a comparison of the running of the mice during the day versus night, was determined. Mean group waveforms were constructed to examine differences in the duration of the active period and rest periods in LD and in DD (period and power values using Clocklab’s Chi Square Periodogram). Period and power of the rhythms were assessed over three blocks in DD (Block one: DD days 1-10, Block two: DD days 48-57, and Block three: DD days 76-84). The difference between the onset of the main nocturnal bout of running and the time of lights off, also known as the phase angle of entrainment (psi), was determined at two points during LD. The duration of the active phase (alpha) and amount of activity, were calculated for the same two points during LD, as well as one point during DD. The effect of masking was determined by compiling the information gathered from the 24 hours the mice spent in the 2:2 LD cycle into 2-hour light and dark periods and then calculating a LD ratio. Additionally, LD ratios were calculated from the masking data by averaging values based on the day and night of the original 12:12 LD cycle. Phase shifts were calculated following exposure to dim (∼40 lux) or bright light (∼1250 lux) at CT16 or CT22 and after exposure to bright light at CT6. Phase shifts were calculated as was described in Experiment 4.

#### Light induced cFos and *mPer1* expression

Two weeks after the completion of the third round of light pulses, cFos and *mPer1* expression following a 15-min bright light pulse was assessed at either CT16 or CT22 in both WT and KO mice (n=3 each genotype and phase). Ninety minutes after exposure to the light pulse, animals were euthanized, in the dark, via an overdose of sodium pentobarbital, and then perfused as described above. Gene expression was also assessed for a second group of WT and KO mice (n=3 each) when administered a bright light pulse at CT22.

#### Image quantification and statistics

Behavioural data for LD ratios, alpha, and psi were analyzed by comparing measures obtained for KO mice with that of WTs. Independent samples t-tests were utilized with significance set at *p*<0.05. Violations of the assumption of homogeneity of variance were assessed using Levene’s test. For power and period data, the effect of time was assessed by using 2×3 mixed factorial ANOVA (genotype: WT, KO x time: block 1, 2, and 3) statistical tests. Phase shifts were analyzed using a 2×2×2 mixed ANOVA (genotype: WT, KO x light: dim, bright x phase: CT16, CT22) with light and phase being the within-subjects factors and genotype the between-subjects factor. As only a bright light pulse was used at the CT6 phase, it was considered in isolation and assessed using an independent samples t-test. In order to compare the expression of the FOS protein and *mPer1* gene in the core versus the shell, 2×2×2 (genotype: WT, KO x phase: CT16, CT22 × region: core, shell) factorial mixed ANOVAs were conducted. Region was the within-subjects factor while genotype and phase were the between-subjects factor. Statistical significance was set at *p*<0.05. All means are reported ± SD in the text and ± SEM in figures.

## RESULTS

### Experiment 1: ZnT3 colocalization with melanopsin retinal ganglion cells

A subset of ipRGCs that contain ZnT3 project to the circadian system (Figure 1). Images of positive staining for melanopsin show the entire ipRGC cell with its large cell body and extensive dendritic arbor visible in the ganglion cell layer. Positive ZnT3 staining was identified as a speckle formation outlining a cell along its perimeter. From a total of 120 ipRGCs, 47.5% were identified as melanopsin single-labelled cells while 52.5% were identified as melanopsin ZnT3 double-labelled cells.

**Figure 1.**
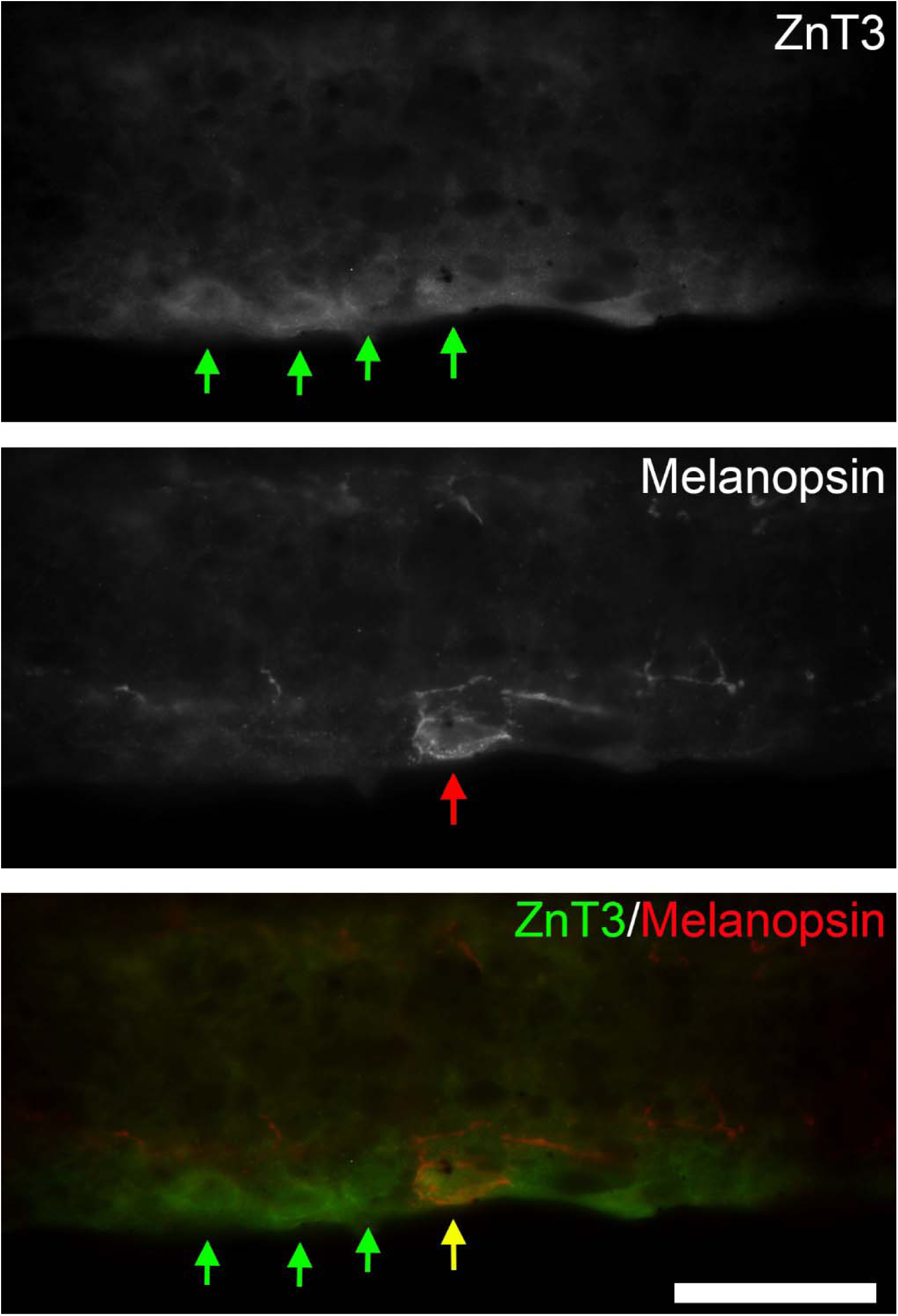
Representative mouse retinal section showing immunofluorescent staining for ZnT3 (top row, green arrows) and melanopsin cells (middle row, red arrow) in the ganglion cell layer (GCL). The bottom row shows a merged image from which colocalization can be deciphered. Three single-labeled ZnT3-IR cells (green arrows) are evident, as is one double-labeled ZnT3+ melanopsin cell (yellow arrow). Scale bar = 30µm.

### Experiment 2: Histochemical zinc expression in the SCN and IGL

A small amount of vesicular zinc staining was present throughout the SCN, which was mostly visible in rostral sections, in the medial and ventral regions (Figure 2A-C). Quantification took place on seven brains, with one animal excluded due to the inability to obtain representative sections. There was no significant difference in vesicular zinc levels (as measured by ROD) between the two time points, ZT6 and ZT18, for the dorsomedial (*t*_(5)_ = 1.879, *p*>0.05), dorsolateral (*t*_(5)_ = 1.634, *p*>0.05) or ventral (*t*_(5)_ =0.875, *p*>0.05) SCN areas (Figure 3A,B). In contrast, a considerable amount of vesicular zinc was present throughout the full rostrocaudal extent of the IGL and extending lower down in the ventrolateral geniculate nucleus (vLGN; Figure 2D-O). Vesicular zinc levels did not significantly differ between ZT6 and ZT18 (*t*_(5)_ =1.518, *p*>0.05, Figure 3C,D).

**Figure 2.**
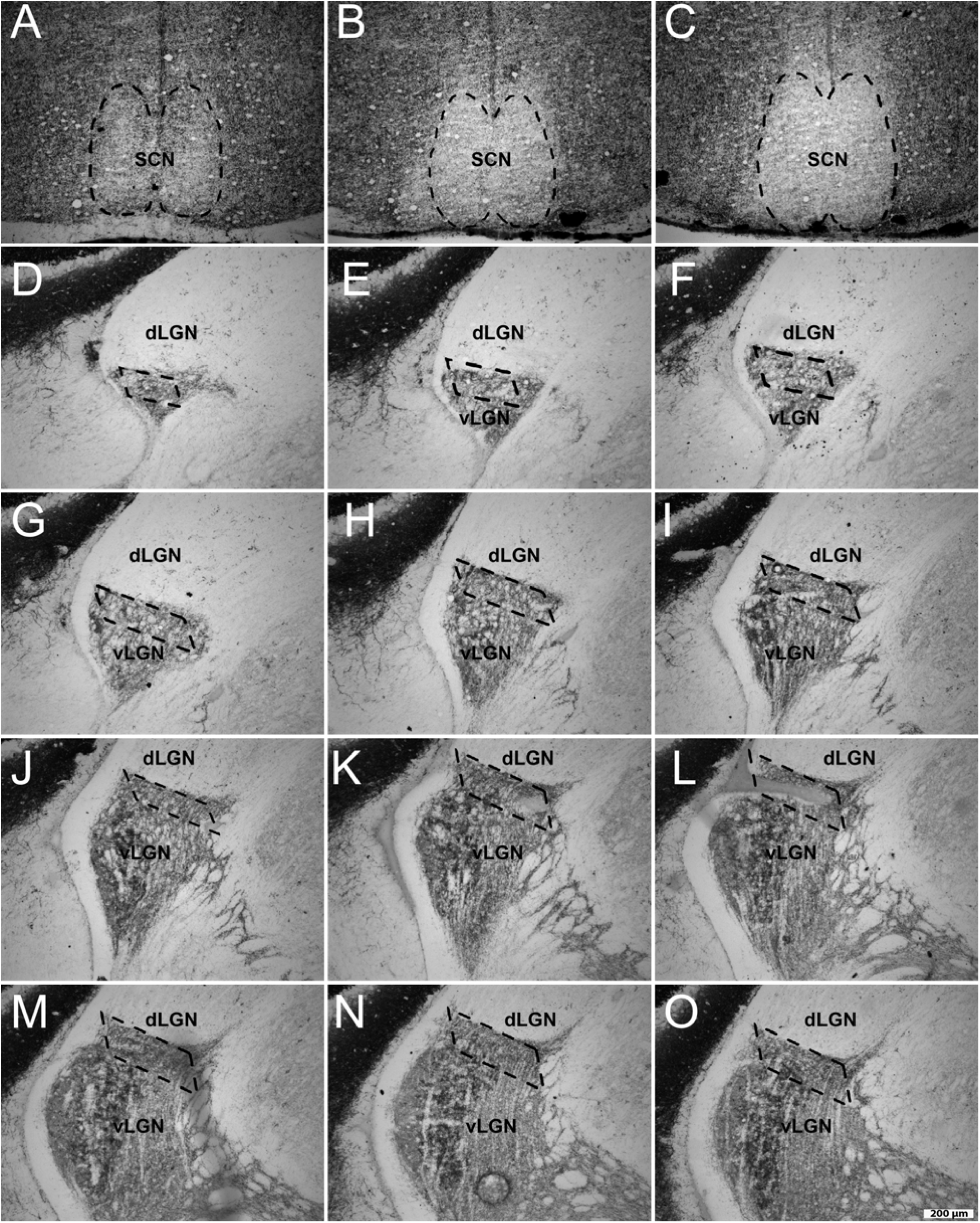
Photomicrographs of representative hamster coronal slices showing autometallographic zinc staining in the suprachiasmatic nucleus (SCN; rostral to caudal direction A-C) and left intergeniculate leaflet (denoted by boxed area; rostral to caudal direction D to O). Vesicular zinc staining extends down into the ventrolateral geniculate nucleus (vLGN) but was completely absent from the dorsolateral geniculate nucleus (dLGN).

**Figure 3.**
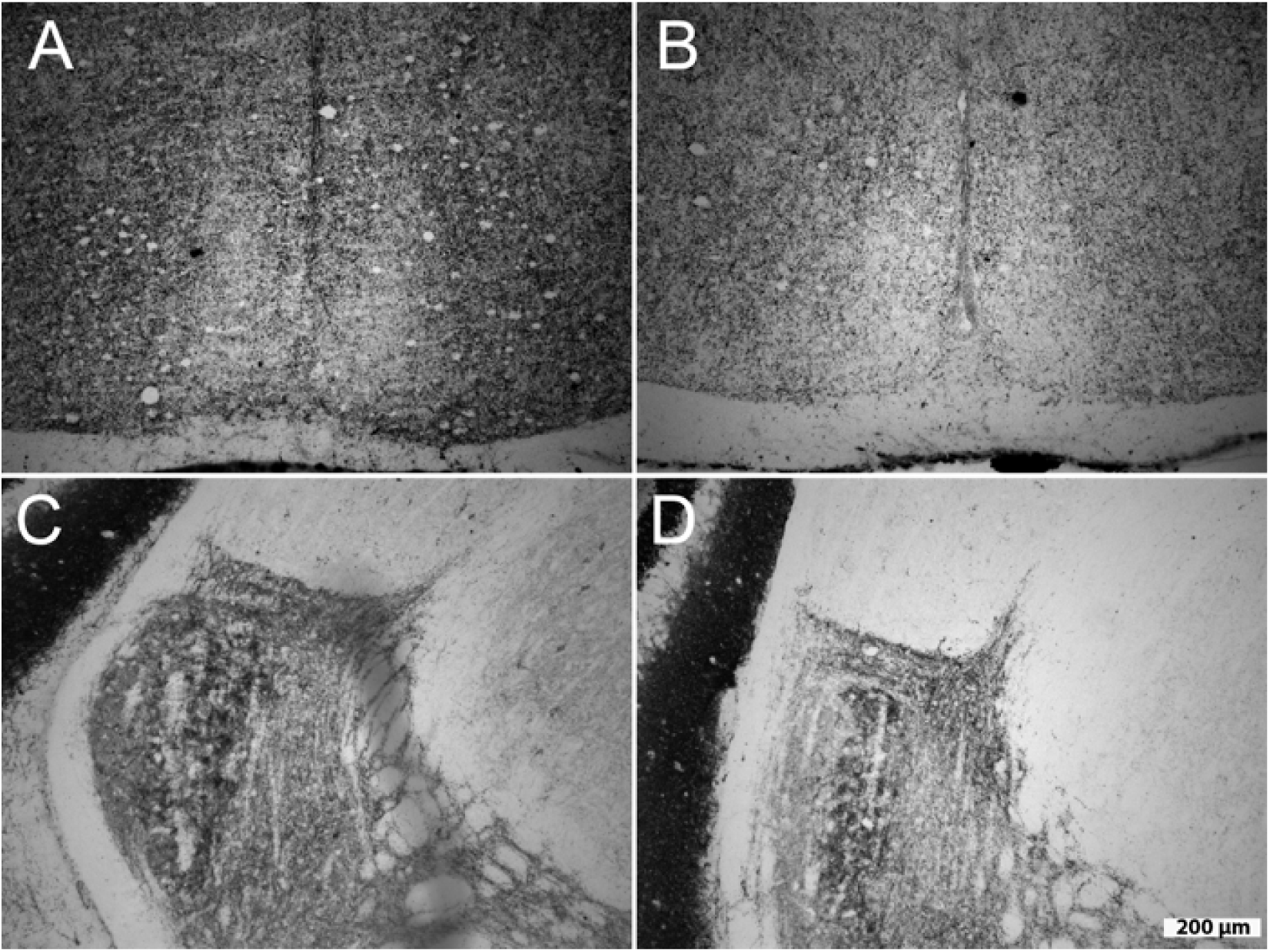
Photomicrographs of representative hamster coronal slices showing levels of vesicular zinc staining in the suprachiasmatic nucleus (SCN, A+B) and the intergeniculate leaflet (IGL, C+D) at ZT6 (A+C) and ZT18 (B+D). No significant differences were found between the time points, *p*>0.05.

### Experiment 3: Vesicular zinc is present in the IGL with or without retinal input

NPY staining was used to delineate the IGL in adjacent sections. Histochemically stained zinc was observed in control animals as well as enucleated animals and there was no significant difference in vesicular zinc staining in the IGL between these two groups (*t*_(4)_ = 1.741, *p*>0.05; Figure 4).

**Figure 4.**
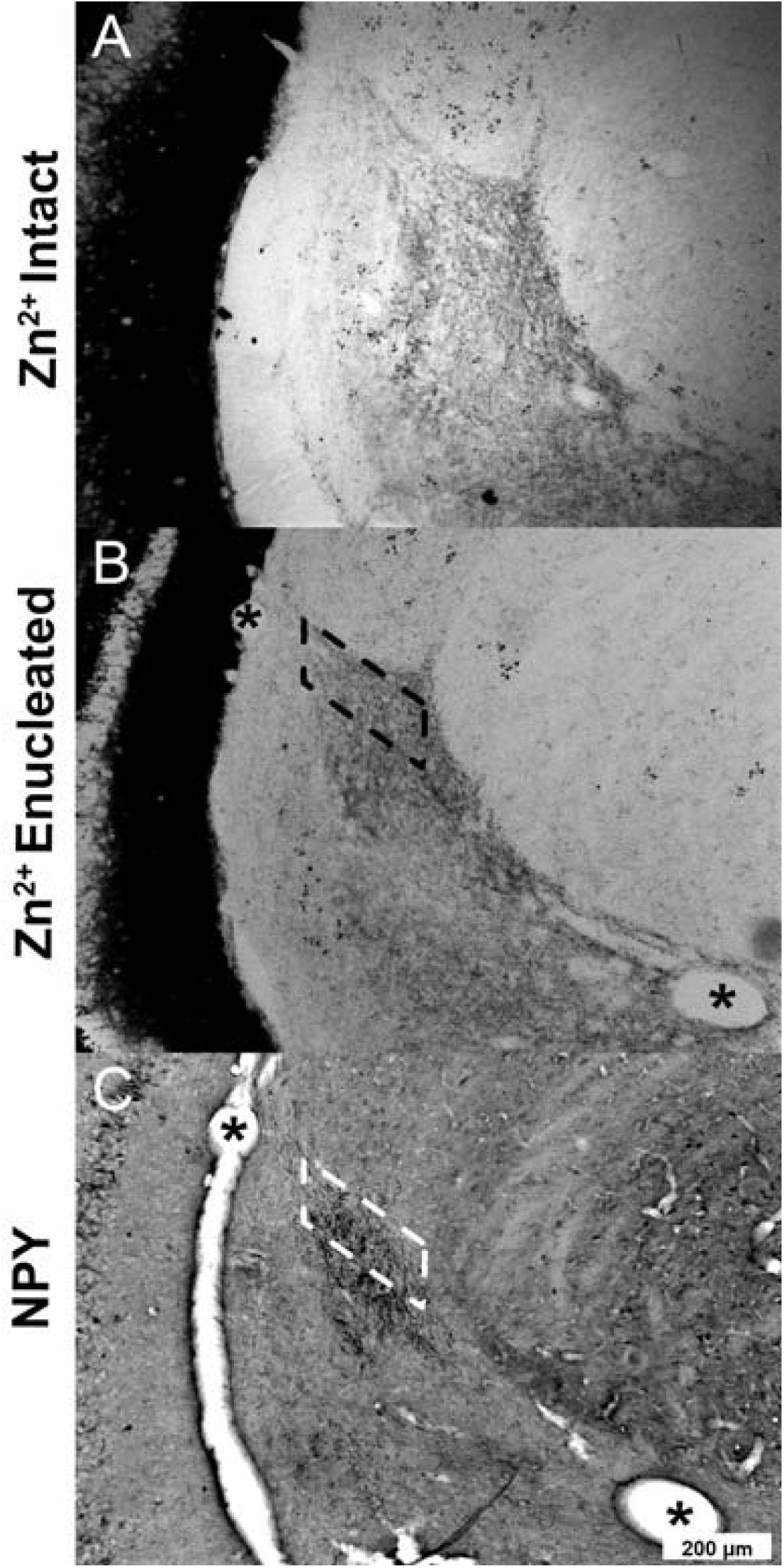
Photomicrographs of representative hamster coronal slices showing vesicular zinc staining in the IGL from animals with intact retinas (A) and from enucleated animals (B). No significant difference was found between the two conditions, *p*>0.05. IGL staining was confirmed by matching an adjacent delineated NPY region (C) to that of the presumed IGL region (B; boxed area). Matching blood vessel landmarks are denoted by (*) for comparison and matching purposes.

### Experiment 4: Zinc level modification at the SCN and IGL does not alter circadian responses to light

All 12 animals were used for analysis for SCN treatments. Data from a total of eight hamsters were used in the analysis for IGL treatments. Histological examination led to three animals being excluded due to missed placements for one of the two cannulas aimed at the left or right IGL. Another animal was excluded after losing its headcap during the first manipulation. Two more animals were excluded due to near zero phase shifts following light exposure in the vehicle control condition. No animals were excluded based on responses to experimental treatments.

Light-induced phase shifts did not differ significantly between the two vehicle treatment conditions of saline and DMSO for either the intra-SCN treatment (saline: 1.820±0.258h; DMSO: 1.686±0.326h; independent t-test, *t*_(10)_ = 0.326, *p*=0.751) or the intra-IGL treatments (saline: 1.854±0.640h; DMSO: 1.924±0.722h; paired t-test, *t*_(12)_ = 0.192, *p*=0.851). As a result, the saline and DMSO data was combined into one vehicle group in these experiments for further analysis with the active treatment conditions (i.e., vs ZnCl_2_ and TPEN conditions). For intra-SCN treatments (Figure 5), there was no significant difference (*F*_(11,22)_=0.926, *p*=0.411) in phase shifts between the vehicle (1.764±0.194h), ZnCl_2_ (2.007±0.239h), or TPEN (1.669±0.193h). Similarly for the intra-IGL experiment, there were no significant differences (*F*_(7,13)_=0.752, *p*=0.491) in phase shifts between the vehicle (1.843±0.432h), ZnCl_2_ (1.619±0.743h), or TPEN (1.571±0.8h) treatments (Figure 6).

**Figure 5.**
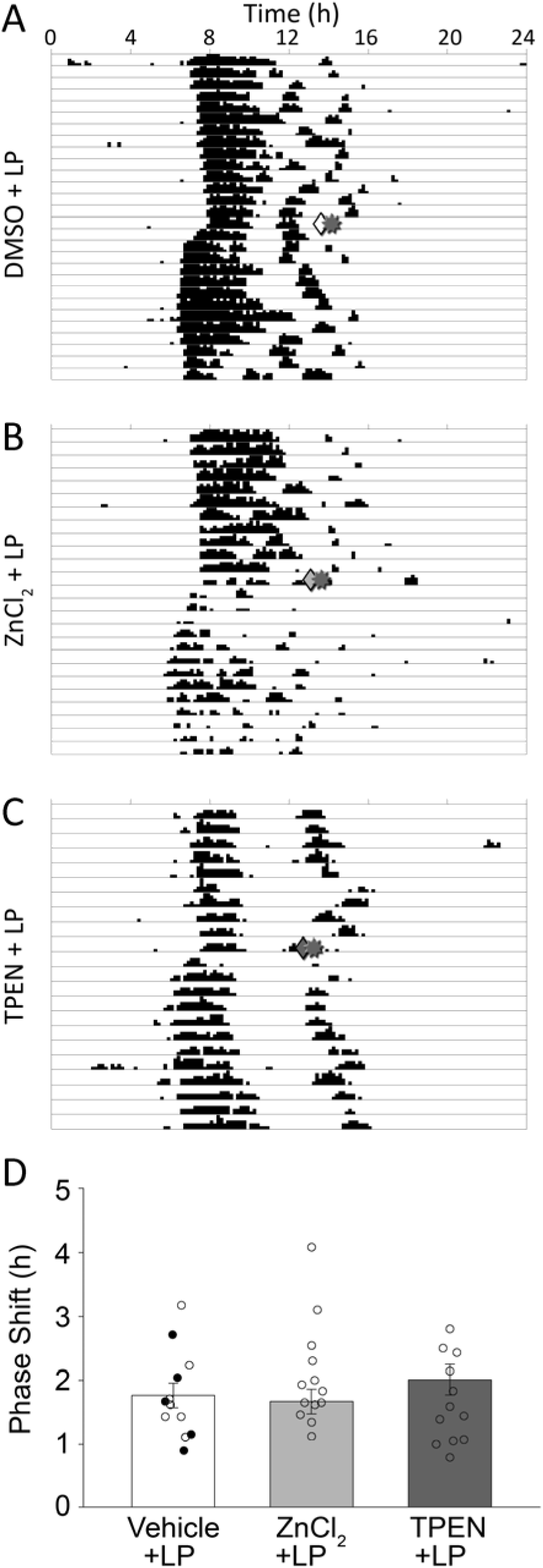
(A-C) Actograms from representative animals depicting the photic phase shift after intra-SCN injection with (A) vehicle (10% DMSO shown) (B) ZnCl_2_ and (C) TPEN, denoted by a diamond (◊) in the SCN10 min prior to a 15 min light pulse (☼; 40lux) at late subjective night (CT18). Each horizontal line represents a day of wheel running as shown by the black vertical bars with subsequent days plotted below. The height of the bars is proportional to the number of wheel revolutions. (D) Mean phase shift (±SEM) of hamsters after treatment with vehicle (average data of saline and DMSO for each animal), ZnCl_2_ and TPEN prior to a 15-minute light pulse (LP) at CT18. For vehicle control, dark circles represent animals receiving 10%DMSO, while open circles represent animals receiving saline. There was no significant difference in photic phase shifts between the treatments, *p*>0.05.

**Figure 6.**
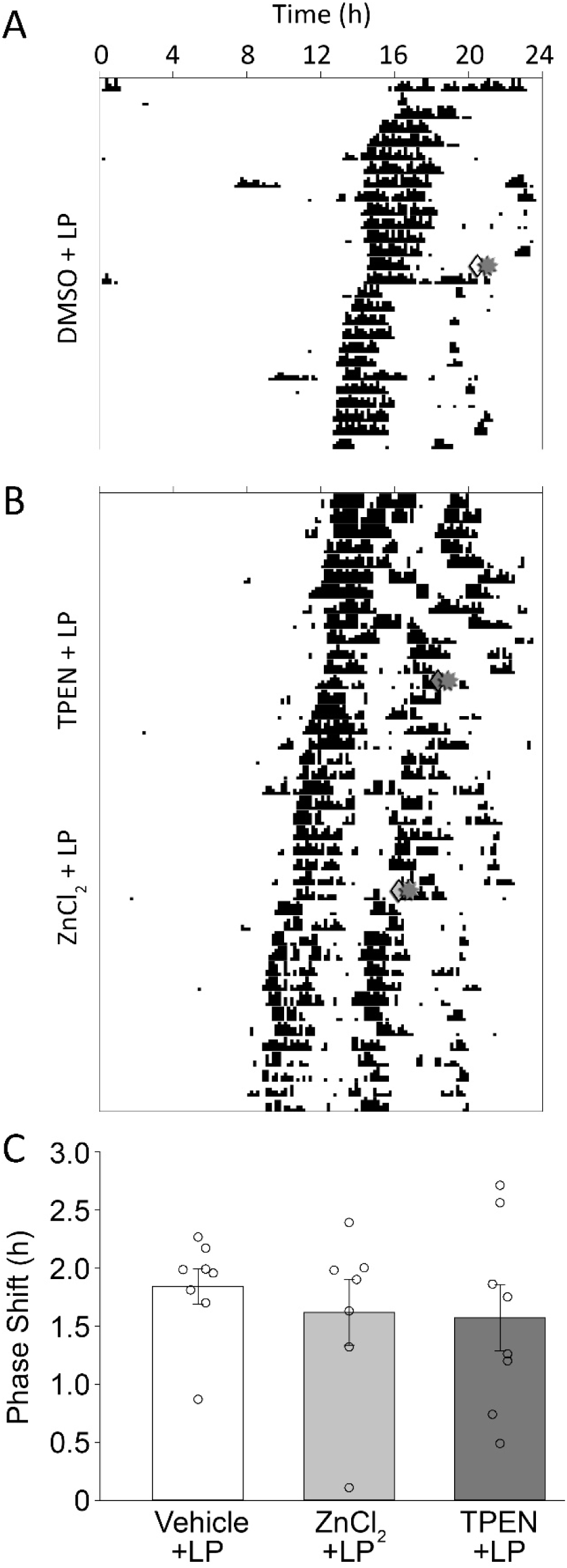
(A-C) Actograms from representative animals depicting the photic phase shift after bilateral intra-IGL injections with (A) vehicle (10% DMSO shown) (B) TPEN and ZnCl_2_, denoted by a diamond (◊) in the SCN10 min prior to a 15 min light pulse (☼; 40lux) at late subjective night (CT18). Each horizontal line represents a day of wheel running as shown by the black vertical bars with subsequent days plotted below. The height of the bars is proportional to the number of wheel revolutions. (C) Mean phase shift (±SEM) of hamsters after treatment with vehicle (average data of saline and DMSO for each animal), ZnCl_2_ and TPEN prior to a 15-minute light pulse (LP) at CT18. There was no significant difference in photic phase shifts between the treatments, *p*>0.05.

### Experiment 5: Circadian rhythms changes in the ZnT3 KO model

#### General circadian characteristics

In a light-dark cycle, ZnT3-KO mice exhibited circadian patterns of wheel running behavior that were similar to wildtype controls. Actograms and average waveforms for the genotypes were of similar appearance (Figure 7). Both genotypes exhibited pronounced nocturnal behavior with limited activity during the light phase. The L/D ratios (i.e., activity in light phase / activity in the dark phase) were not significantly different between the genotypes (WT: 0.029±0.022; KO: 0.036±0.028; *t*_(10)_=0.474, *p*=0.646). The duration of the active phase (alpha) was similar between the genotypes (LD: WT=6.35±0.8h, KO=5.8±1.0h; *t*_(10)_=1.02, *p*=0.331; DD: WT=6.6±1.0h, KO=5.8±1.59h, *t*_(10)_=1.046, *p*=0.320). Both wildtype and ZnT3-KO mice exhibited small positive phase-angles of entrainment (WT: 6.4±0.8 min, KO: 5.8±1.0 min) that were not significantly different (*t*_(10)_=1.02, *p*=.331).

**Figure 7.**
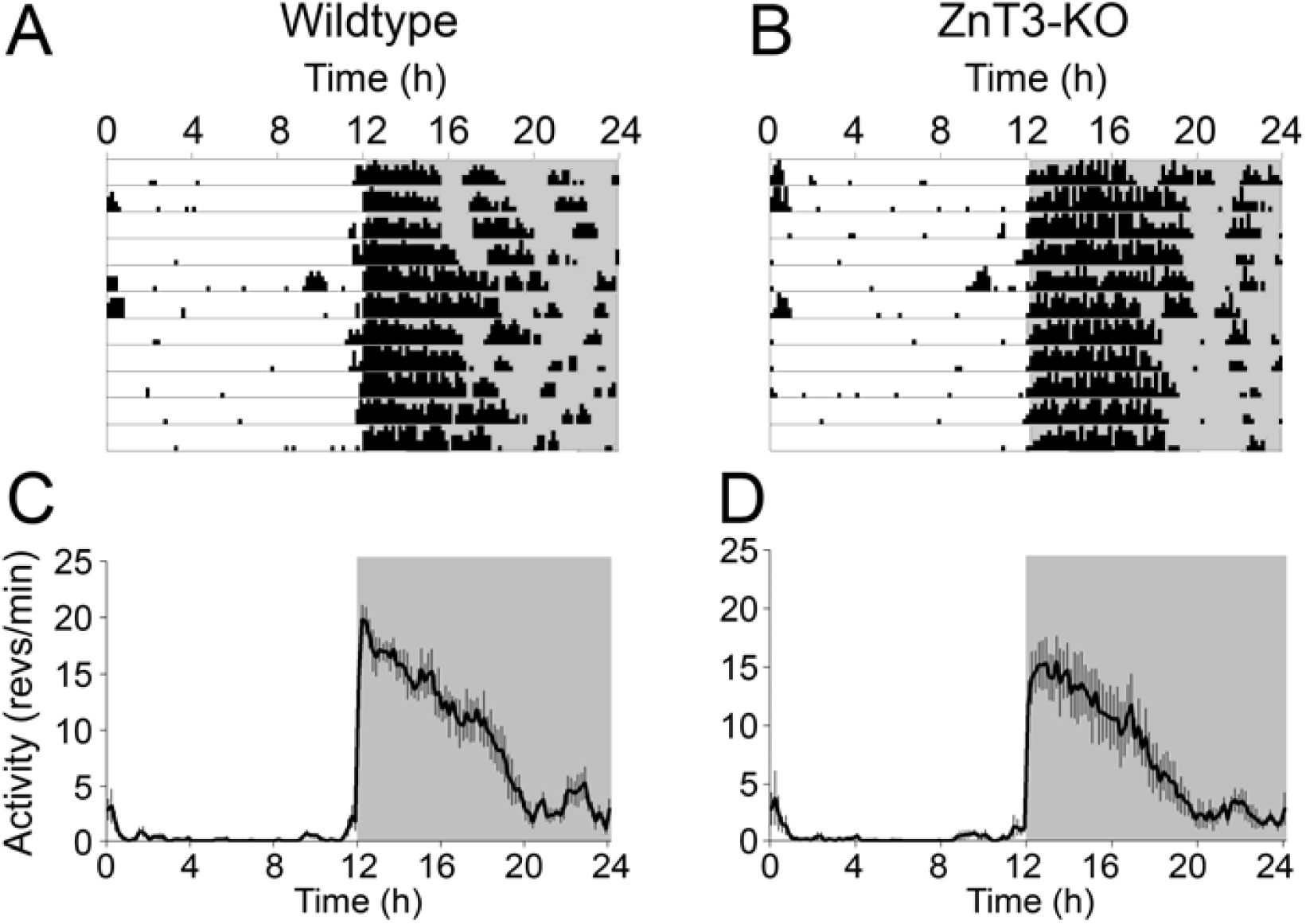
Representative actograms from wildtype (A) and ZnT3-KO (B) mice in an LD cycle. Average waveforms of wheel running activity from all wildtype (C) and ZnT3-KO (D) mice over the days depicted in the actograms. The vertical gray bars represent the SEM across animals.

In constant darkness, both WT and KO mice exhibited clear free-running rhythms (Figure 8) that did not significantly differ in terms of their period (WT= 23.67±0.1h, KO=23.64±0.2h; *F*_(1,10)_=0.090, *p*=0.770) or their power (Max χ^2^ power: WT=716.19±228.19, KO=771.73±260.62; *F*_(1,10)_=0.586, *p*=0.462). Period and power significantly decreased with time in DD (main effect of block for both period, *F*_(2,10)_=15.06, *p*<0.001, and power, *F*_(2,10)_=31.32, *p*<0.001), but this change was similar for both genotypes (genotype X block interaction for period, *F*_(2,10)_=0.595, *p*=0.561, and power, *F*_(2,10)_=0.015, *p*=0.985).

**Figure 8.**
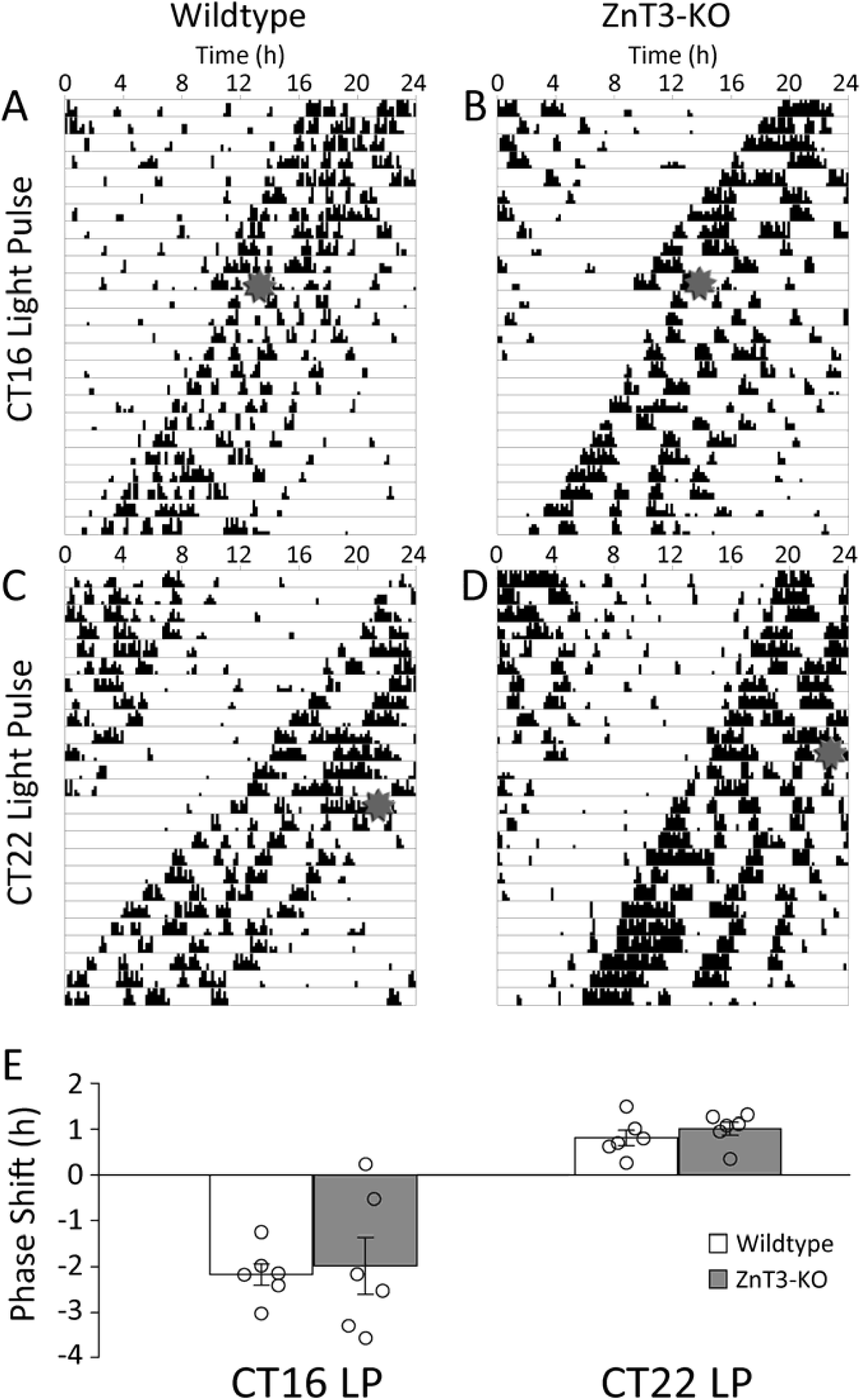
Representative actograms from wildtype (A,C) and ZnT3-KO (B,D) mice depicting phase shifts to a 15 min, 1250lux light pulse at CT16 (A,B) and CT22 (C,D). E) Mean shifts (±SEM) from wildtype mice (white bars) and ZnT3-KO mice (gray bars). and individual shifts (circles). No significant difference in phase shifts were observed between the genotypes at either circadian phase examined.

#### Masking

For the masking paradigm, LD ratios (see above) were calculated for activity that occurred during the 12h day, 12h night, and the full 24h period when animals were housed in a LD 2:2 cycle. During the day, the ratio of activity when the lights were on compared to when the lights were off was not significantly different between the WT mice (0.038±0.026) and KO mice (0.252±0.430; *t*_(10)_=1.22, *p*=0.251). Similarly, during the night, LD ratios were not significantly different between WT mice (0.060±0.030), and KO mice (0.098±0.106; *t*_(10)_=0.842, *p*=0.419). Total LD ratios were also not significantly different between WT (0.058±0.028) and KO mice (0.100±0.101; *t*_(10)_=0.975, *p*=0.352).

#### Phase shifts

Phase shifts were examined following exposure to dim (40lux) and bright (1250lux) light at CT16 and CT22, as well as following exposure to bright light at CT6. Representative phase shifts to the bright light at CT16 and CT22 are displayed in Figure 9. Large phase delays and advances were observed to light pulses at CT16 and CT22 respectively. At both CT16 and CT22 phase shifts were significantly larger with bright light exposure (CT16 *F*_(1,10)_=5.384, *p*=0.043; CT22 *F*_(1,10)_=28.884, *p*<0.001). In neither case were the phase shifts significantly different between the genotypes (CT16 *F*_(1,10)_=0.552, *p*=0.475; CT22 *F*_(1,10)_=0.271, *p*=0.614). There were no significant interactions between genotype and light intensity (CT16 *F*_(1,10)_=0.00176, *p*=0.967; CT22 *F*_(1,10)_=0.325, *p*=0.581). Small phase delays were observed to bright light pulses at CT6 but did not significantly differ between the genotypes (WT −0.37±0.532h, KO −0.578±0.385, *t*_(9)_=0.762, *p*=0.466).

**Figure 9.**
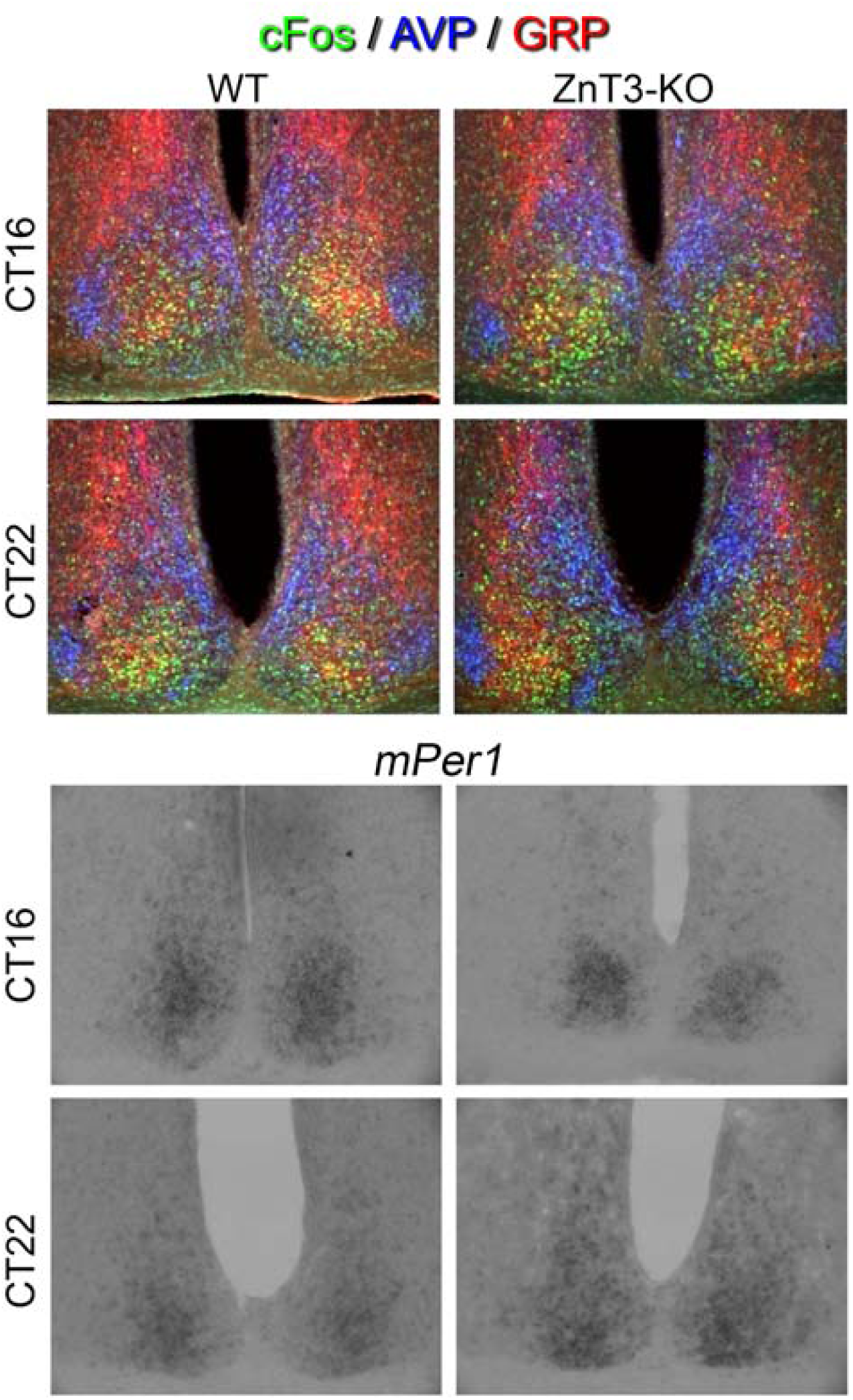
Representative photomicrographs from wildtype (WT, left column) and ZnT3-KO mice (right column) depicting light induced cFos (top set) and *mper1*(bottom set) from animals given a 15 minute, 1250lux light pulse at CT16 (top row of each set) or CT22 (bottom row of each set). Immunoreactivity for arginine vasopressin (AVP, blue) was used to delineate the SCN shell, while immunoreactivity for gastrin-releasing peptide (GRP, red) was used to delineate the SCN core when counting cells with cFos immunoreactive nuclei (green). No significant differences were detected between the genotype for cFos or *mper1* expression at either circadian phase.

#### Light-Induced Gene Expression

Light pulses (15min, 1250lux) induced prominent Fos expression in the SCN (Figure 9). Regardless of phase or genotype, greater FOS expression was observed in the SCN core than in the SCN shell (*F*_(1,8)_=148.87, *p*<0.001). None of the interactions involving region were significant (phase: *F*_(1,8)_=0.611, *p*=.457; genotype: *F*_(1,8)_=0.296, *p*=0.601; phase x genotype: *F*_(1,8)_=0.235, *p*=0.651). Additionally, no significant differences in FOS expression were observed in relation to phase (*F*_(1,8)_=1.09, *p*=0.326) or genotype (*F*_(1,8)_=0.007, *p*=0.933). The interaction between phase and genotype was also not significant (*F*_(1,8)_=0.002, *p*=0.964).

Light pulses also induced prominent *mPer1* expression in the SCN (Figure 9). Regardless of phase or genotype, greater *mPer1* expression was observed in the SCN core than in the SCN shell (*F*_(1,14)_=89.88, *p*<0.001). No significant differences in *mPer1* expression were observed in relation to phase (*F*_(1,14)_=0.191, *p*=0.668) or genotype (*F*_(1,14)_=0.075, *p*=0.789). The interaction between region and phase was not significant (*F*_(1,14)_=0.007, *p*=0.932) nor was the interaction between region and genotype, *F*_(1,14)_=0.349, *p*=0.564. The three-way interaction between region, phase, and genotype was also not significant, *F*_(1,14)_=1.40, *p*=0.256.

## DISCUSSION

The present study is the first to thoroughly examine the anatomical distribution and functional role of vesicular zinc in the circadian system. Consistent with previous reports (Redenti and Chappell, 2004), we localized the vesicular zinc transporter ZnT3 to the ganglion cell layer of the retina, and demonstrated that a subset of melanopsin-containing ipRGCs colocalized ZnT3. Minimal vesicular zinc was observed in the hamster SCN, contrasting with previous reports that detected zinc in the rat SCN (Huang, et al., 1993). However, strong histochemically-stained zinc was detected throughout the entire IGL structure. Vesicular zinc staining was also observed throughout the ventrolateral geniculate. Zinc staining levels in the SCN and IGL did not change across the circadian day. While retinal ganglion cells project to the lateral geniculate and IGL, loss of the eyes did not significantly alter vesicular zinc staining in these structures, suggesting that they receive significant zinc input from elsewhere in addition to or instead of from the retina. While some ipRGCs were found to colocalize ZnT3, allowing them to synaptically release zinc, increasing or decreasing zinc in either the SCN or IGL failed to significantly alter circadian responses to phase shifting light pulses in hamsters. Similarly, mice lacking ZnT3 which have no vesicular zinc, did not exhibit any differences from wildtype mice in entrained, free-running or phase shifting circadian responses. It is possible that other circadian tasks, such as anticipation of scheduled feeding, or other non-photic manipulations, might reveal a difference between the WT and ZnT3 mice.

The colocalization of ZnT3 with ipRGCs suggest a possible function of vesicular zinc as a modulator in the brain areas innervated by ipRGCs. A number of ipRGC subtypes have been noted, with distinct morphologies and projection patterns (Berson et al., 2010;Sexton et al., 2012;Schmidt et al., 2011). Specifically, the M1 type lacking BRN3b project to the SCN while the M2 type that contains BRN3b project to the IGL. Many of the melanopsin-positive cells that were observed to contain ZnT3 had their dendrite more in the ON sublamina of the inner plexiform layer, suggesting that these were M2, M3, M4 or M5 ipRGCs, which project to structures such as the dorsolateral geniculate, the superior colliculus, or the core of the olivary pretectal nucleus (Schmidt et al., 2011). The lack of change of zinc staining in the IGL and vLGN of enucleated hamsters does not exclude the possibility that the retina provides zincergic input, but at a minimum requires that there also be other significant sources of vesicular zinc input. The IGL receives considerable afferent projections from several different brain areas including visual centers such as the superior colliculus (SC), visual cortex and dLGN (Moore et al., 2000), the hypothalamus, and the raphe nucleus. Of these, the visual cortex is known to contain zincergic neurons (Brown and Dyck, 2004). While manipulations of zinc levels in the hamster IGL, or globally with the ZnT3-KO mouse failed to alter circadian responses to light, the IGL is also known to participate in non-photic phase shifting (Webb et al., 2014). It is possible that zinc may play a role in regulating these non-photic responses. Photic stimuli interfere with non-photic responses (Mrosovsky, 1991) so it is possible that zincergic input to the IGL might modulate this phenomenon.

Vesicular zinc is found in high levels in the retina, largely in the photoreceptor cells, amacrine cells, and retinal pigment epithelial cells (Ugarte and Osborne, 2001). The location of vesicular zinc in photoreceptors changes with the light cycle, but this appears to be a light-regulated rather than a circadian-regulated feature (Ugarte and Osborne, 2001). In the retina, vesicular zinc may play a number of roles. ZnT3 is found in a number of retinal cells (Redenti and Chappell, 2004), and synaptically-released zinc appears to modulate NMDA and GABA receptors in a number of retinal circuits (Ripps and Chappell, 2014;Ugarte and Osborne, 2001). Zinc also regulates the activity of retinol dehydrogenase necessary for the production of retinal. This latter role is thought to explain night-blindness and poor dark adaptation in people with zinc deficiencies (Ugarte and Osborne, 2001). Patients with zinc deficiencies take longer to adapt to darkness and are less sensitive when fully adapted (Morrison et al., 1978). Circadian responses to light are influenced by dark adaptation, with responses to light increasing with time in constant darkness (Daymude and Refinetti, 1999;Refinetti, 2001;Shimomura and Menaker, 1994). However, the timescale for maximal dark adaptation of circadian responses, about 3 weeks, far exceeds the timescale for dark adaptation of visual responses in people (∼40 min;Pirenne, 1962). Given the lack of difference in phase shifts between WT and ZnT3-KO mice, it is likely that if vesicular zinc plays a role in dark adaptation of the circadian response, then this many be mediated by free zinc rather than synaptically-released zinc that requires the ZnT3 transporter.

A role for vesicular zinc in the SCN has been suggested previously (Huang, et al., 1993). In rats, using a technique similar to that employed here, those authors detected prominent zincergic input to the ventral region of the SCN. It was found that zinc enhanced inhibitory outward potassium I_A_ currents in isolated SCN neurons from the rat (Huang, et al., 1993). GABA_A_ currents are also zinc sensitive in the rat SCN (Kawahara, et al., 1993;Strecker, et al., 1999). The sensitivity of these GABA-induced current to zinc varies over the day (Kretschmannova et al., 2005;Kretschmannova et al., 2003). Finally, NMDA currents in the SCN are also inhibited *in vitro* by zinc, an observation which has been used as evidence of the presence of the zinc-sensitive GluN2A subunit of the NMDA receptor (Clark and Kofuji, 2010). The lack of responses to manipulations of zinc in our study could reflect species differences between rats used in these other studies and hamsters and mice explored here. Alternatively, the pharmacological responses observed *in vitro* in these other studies may not translate to the intact behaving animals explored here.

The transgenic mouse model used here has lacked the ZnT3 transporter since fertilization, thus there may be developmental compensation to mitigate the persistent lack of vesicular zinc. Similar to what we observed with our circadian investigations, ZnT3 mice do not differ from WT mice on a wide variety of behavioural tasks (Cole et al., 2001). Recent evidence suggest that differences do start to emerge when tasks are designed to be especially sensitive/challenging, or when the animals are subjected to stress (McAllister et al., 2018;Thackray et al., 2017;Wu and Dyck, 2018).

Zinc is a critical essential trace element that participates in a wide variety of biochemical processes. With the present study, we have found that the transporter responsible for packaging zinc into synaptic vesicles is located in a subset of melanopsin-containing ipRGCs, and that vesicular zinc is found throughout the IGL. Despite this, altering zinc levels in the circadian system with zinc donors or chelators did not alter circadian responses to light, and genetic deletion of the necessary transporter for packaging of vesicular zinc did not alter any of the entrained, free-running or phase shifting circadian responses. While our study provides evidence for the presence of vesicular zinc in the wider circadian network (i.e., retina and IGL), our current approaches failed to find a functional role for vesicular zinc in the circadian system.

## Acknowledgements

This work was supported by Natural Sciences and Engineering Research Council of Canada (NSERC) Discovery Grants to RHD and MCA and by a Canadian Institutes of Health Research (CIHR) Operating Grant to RHD.

## Abbreviations

ANOVA: Analysis of variance
AVP: Arginine vasopressin
CT: Circadian time
DD: Constant darkness
dLGN: Dorsolateral geniculate nucleus
DMSO: Dimethyl sulfoxide
GABA: γ-aminobutyric acid
GCL: Ganglion cell layer
GHT: Geniculohypothalamic tract
GRP: Gastrin-releasing peptide
I_A_: Transient potassium current
IGL: Intergeniculate leaflet
INL: Inner nuclear layer
i.p.: Intraperitoneal
IR: Immunoreactive
IPL: Inner plexiform layer
ipRGCs: Intrinsically photosensitive retinal ganglion cells KO Knockout
LD: Light-dark cycle
LP: Light pulse
NMDAR: N-methyl-D-aspartate receptor
NPY: Neuropeptide Y
OPL: Outer plexiform layer
PBS: Phosphate buffered saline
PBSx: PBS with Triton-X-100
PCR: Polymerase chain reaction
PFA: Paraformaldehyde
psi: Phase angle of entrainment
SCN: Suprachiasmatic nucleus
SD: Standard deviation
SEM: Standard error of the mean
TPEN: N,N,N’,N-Tetrakis(2-pyridylmethyl) ethylenediamine
VGLUT1: Vesicular glutamate transporter 1
VIP: Vasoactive intestinal polypeptide
vLGN: Ventrolateral geniculate nucleus
WT: Wildtype
ZIP: ZRT/IRT-like proteins
ZnCl_2_: Zinc Chloride
ZnT: Zinc transporter
ZnT3: Zinc transporter 3
ZT: Zeitgeber time

